# Error-based implicit learning in language: the effect of sentence context and constraint in a repetition paradigm

**DOI:** 10.1101/2023.12.13.571412

**Authors:** Alice Hodapp, Milena Rabovsky

**Affiliations:** University of Potsdam, Potsdam, Germany

## Abstract

Prediction errors drive implicit learning in language, but the specific mechanisms underlying these effects remain debated. This issue was addressed in an electroencephalogram (EEG) study manipulating the context of a repeated unpredictable word (repetition of the complete sentence or repetition of the word in a new sentence context) and sentence constraint. For the manipulation of sentence constraint, unexpected words were presented either in high constraint (eliciting a precise prediction) or low constraint sentences (not eliciting any specific prediction). Repetition induced reduction of N400 amplitudes and of power in the alpha/beta frequency band was larger for words repeated with their sentence context as compared to words repeated in a new low constraint context, suggesting that implicit learning happens not only at the level of individual items but additionally improves sentence-based predictions. These processing benefits for repeated sentences did not differ between constraint conditions, suggesting that sentence-based prediction update might be proportional to the amount of unpredicted semantic information, rather than to the precision of the prediction that was violated. Additionally, the consequences of high constraint prediction violations, as reflected in a frontal positivity and increased theta band power, were reduced with repetition. Overall, our findings suggest a powerful and specific adaptation mechanism that allows the language system to quickly adapt its predictions when unexpected semantic information is processed, irrespective of sentence constraint, and to reduce potential costs of strong predictions that were violated.

## Introduction

Our past sensory experiences shape future perception: Neural computations have been suggested to rely on an internal model of the statistics of our environment. Based on these statistics, predictions of upcoming input are generated (e.g., Elman, 1990; Friston, 2005; McClelland, 1994; Rao & Ballard, 1999; Schultz et al., 1997). When these predictions are disconfirmed, the resulting error signals are thought to promote implicit learning (den Ouden et al., 2009; Elman, 1990; Friston, 2005; McClelland, 1994; McLaren, 1989; Rao & Ballard, 1999; Schultz & Dickinson, 2000), which enables adaptation to new situations with the aim to make better predictions in the future. Our language system can draw on vast experience over the course of our lives and it’s becoming increasingly accepted that it uses this experience to implicitly predict upcoming information and continuously adapts to improve these predictions (Bornkessel-Schlesewsky & Schlesewsky, 2019; Fitz & Chang, 2019; Kuperberg et al., 2020; Rabovsky et al., 2018), while the details of this error-based learning mechanism are still up to debate.

### Mechanisms underlying N400 repetition effects

One electrophysiological correlate of predictive processing in language is the N400 ERP component, a centro-parietal negative going brain potential that peaks around 300-500 ms after word onset and is gradually reduced with higher word predictability (Kutas & Hillyard, 1980, 1984). Given this high sensitivity to predictability, the N400 is often interpreted as an index of unpredicted semantic information (e.g., Bornkessel-Schlesewsky & Schlesewsky, 2019; Federmeier et al., 2007; Kuperberg et al., 2020; Lindborg et al., 2023; Rabovsky et al., 2018). The N400 amplitude becomes smaller when words are repeated (e.g., in word lists: Rugg, 1985). Crucially, word predictability influences the repetition effect. Repeating a semantically incongruent (and thus unpredicted) sentence continuation leads to a long lasting and stronger N400 repetition effect (i.e., a repetition induced reduction of N400 amplitudes), as compared to repeating previously predictable sentences (Besson et al., 1992; Lai et al., 2021; Mitchell et al., 1993; Rommers & Federmeier, 2018b, 2018a). As the N400 component is sensitive to unpredicted semantic information, a reduced amplitude for repeated information can thus be interpreted as an index of changed predictions. Together, this implies that there is a stronger update of predictions for previously unpredictable compared to predictable input.

Interestingly, this larger N400 amplitude reduction for previously unexpected information with repetition has been separately demonstrated for sentence repetitions (Besson et al., 1992) and words repeated in a new sentence context (Rommers & Federmeier, 2018b). This raises questions about the underlying mechanism(s). Error-based learning accounts of language generally assume that unexpected input is processed more thoroughly (Rommers & Federmeier, 2018b) and will update probability representations making the unexpected word less surprising if it is encountered again (Hodapp & Rabovsky, 2021; Rabovsky et al., 2018; Rabovsky & McRae, 2014). Previous research findings on repeated presentation of sentences as well as words in new sentences could therefore each be explained by changes to the representation of the unexpected words themselves. However, most accounts of prediction in language processing (implicitly or explicitly) assume a mechanism that goes beyond such word-level memory or update effects: prediction errors should not only update the representation of the unexpected word but also the predictions made from the respective context, i.e., the connections between the context and the previously unexpected word (explicitly modeled in Rabovsky et al., 2018). Expecting previously surprising input to appear again is useful, but it would be even more effective to know in which context to predict this information. Here, we want to investigate this assumption by comparing N400 amplitudes of words that are repeated in the same sentence to those that are repeated in a new context. When a whole sentence is repeated within the experiment it can (additionally) benefit from updated predictive connections between the word and its context, while results from a previously seen word, that is repeated in a new sentence, can only reflect word-level adaptive processes. Studies that used explicit instructions to memorize items and/or had a behavioral memory test, showed first results that suggest differences between these types of repetition (Besson & Kutas, 1993; Mitchell et al., 1993). For the question raised however, it is crucial that any learning happens incidentally and is not task relevant. In summary, the first experimental manipulation focused on the levels at which unexpected input influences future predictive language processing (this comparison is from now on referred to as context manipulation). If upon processing unexpected information, our predictive language system adapts its predictions made from sentence context beyond adaptation to the unexpected word itself, we would expect to see a stronger N400 amplitude reduction (from first to second presentation) for repeated sentences as compared to repetitions of the same target word in a new sentence context (Fig. 1A).

**Figure 1.**
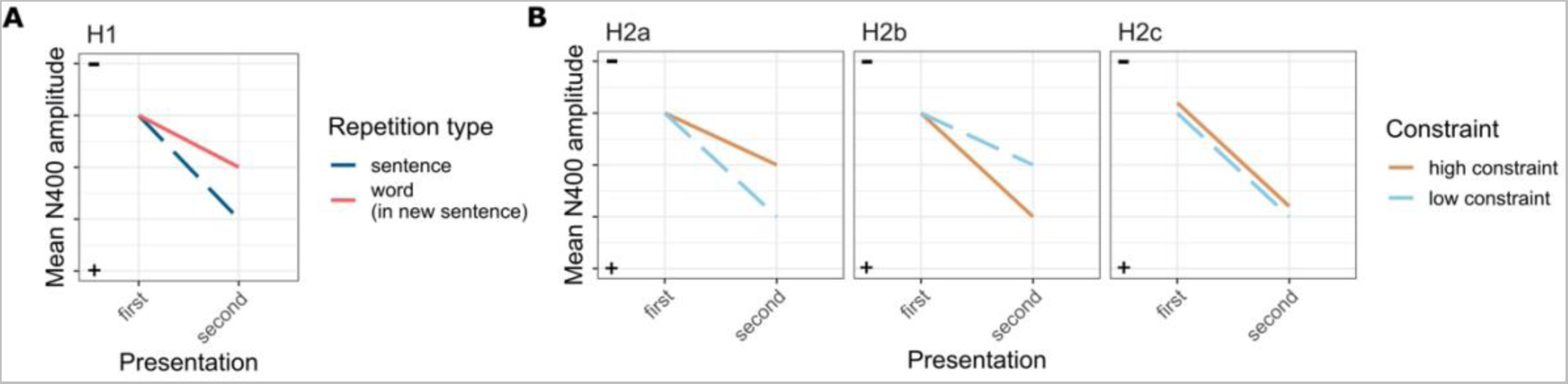
Negative is plotted upwards in line with ERP plots. (A) Hypothesis for the context manipulation (B) Different hypotheses for the constraint manipulation. Conditions in H2c have a slight offset for visualization only and do not reflect a hypothesized difference in repetition effect.

The N400 to unexpected input is invariant to sentential constraint (Federmeier et al., 2007; Kutas & Hillyard, 1984), which is operationalized as the cloze probability of the most frequent response in a sentence completion task. The comparable N400 amplitudes are in line with the equal amount of unpredicted semantic information induced by unexpected continuations (or equal cloze probabilities for the unexpected word), regardless of the sentence’s constraint. However, in strongly constraining sentences, the most frequent response will have a high cloze probability, meaning that any other continuation will violate a precise prediction (e.g., “The children went outside to *look*” when “play” would have been expected). Since low constraint sentences are (by definition) not predictive of a specific continuation, any continuation will be unexpected but will not violate any specific expectations (e.g., “Joy was too frightened to *look*”; both example sentences taken from Federmeier et al., 2007). Despite these differences, unexpected words that were encountered in a highly constraining or weakly constraining sentence elicited not only a similar N400 at initial presentation, but they also showed a comparable more positive N400 if the word was presented again in a new sentence (Lai et al., 2021), indicating that the adaptation processes to the unexpected word itself are not influenced by sentence constraint. The hypothesis that repeated sentences capture additional cognitive processes compared to repeated words (context manipulation, see above), raises the follow-up question of how constraint of sentence context, and therefore the precision of the prediction that was violated by the unexpected information, influences future predictions made from this context. While unexpected information is expected to induce adaptation in high as well as low constraint sentences (Besson et al., 1992; Besson & Kutas, 1993; Lai et al., 2021), their repetition effects have, to the best of our knowledge, not been compared. Previous literature suggests three possible ways in which prediction strength might modulate the rate of this adaptation (Fig. 1B).

On the one hand, repetition effects for unexpected words in high constraint sentences might be reduced compared to repetitions of unexpected words in low constraint sentences. It intuitively seems maladaptive to update representations after an experience that was highly unlikely based on previous experience, so taking the certainty of the prediction into account could keep the internal model from over-adapting to the potentially noisy input. It has been shown that previously expected words that were not encountered nevertheless show a reduced N400 amplitude if presented later (Rommers & Federmeier, 2018a). These ‘lingering expectations’ seem to have no influence on the repetition effects of the unexpected words themselves (Lai et al., 2021), but if the prediction is strong enough to elicit a memory trace even with the prediction disconfirmed (Foucart et al., 2015; Hartman & Hasher, 1991; Hubbard et al., 2019; Rommers & Federmeier, 2018a), this false memory could interfere when the sentence context is repeated. Specifically, surprising continuations in highly constraining sentences would not only elicit an implicit memory of the overall sentence with the unexpected continuation but also of the (not seen) expected item. This could reduce the prediction update, if compared to surprising continuations in weakly constraining sentences for which only one implicit representation of the processed sentence meaning exists (see hypothesis H2a in Fig. 1B).

On the other hand, it is possible that high precision predictions that are violated (i.e., high constraint unexpected sentences) increase prediction update by indicating a change in the environment (unexpected uncertainty: e.g., O’Reilly, 2013; Yu & Dayan, 2005).

Furthermore, in declarative learning the precision of the prior seems to be an important determinant for downstream memory performance. Novel word learning benefits from highly constraining contexts (Gambi et al., 2021) and while less expected words are generally associated with better recognition memory (Corley et al., 2007; Federmeier et al., 2007) and free recall (McFalls & Schwanenflugel, 2002), this effect seems to be strongest for unexpected words that violate a precise prediction (Federmeier et al., 2007). This overcorrection effect for strong predictions is known in domains other than language (Butterfield & Metcalfe, 2006) and could prevent possible costs of predictions gone wrong (hypothesis H2b in Fig. 1B).

Lastly, it is possible that prediction update after unexpected sentence continuations is not sensitive to sentence constraint. Work that explicitly modeled prediction error-based updates of connection weights in a computational model (Rabovsky et al. 2018), updated these connections in proportion to the amount of unpredicted semantic information as reflected in the N400 amplitude. This thus raises the possibility that the update of sentence-based predictions – just like word-level update – is only proportional to the amount of unpredicted semantic information processed and is neither reduced nor enhanced by the precision of violated predictions (hypothesis H2c in Fig. 1B).

In sum, the second experimental manipulation investigated the role of sentence constraint on downstream predictive processing (this comparison is from now on referred to as constraint manipulation). For this, the current study compared high constraint unexpected sentences (which inherently violate precise predictions) to low constraint unexpected sentences (unexpected input due to a less predictive context, but not violating specific predictions) and examined the reduction of N400 amplitude from first to second presentation of the sentence. If the combination of high confidence predictions and unexpected input leads to reduced adaptation, we would expect to see a smaller amplitude reduction for high constraint compared to low constraint sentences (H2a in Fig. 1B). On the other hand, adaptation might increase with the precision of the violated prediction. This would be reflected in a stronger amplitude reduction for the high constraint compared to the low constraint condition (H2b in Fig. 1B). If, however, the overall language model update relied on unpredicted semantic information rather than certainty (and therefore the strength of violated predictions), we would expect to find the same level of N400 amplitude reduction for both conditions (H2c in Fig. 1B).

### Post-N400 repetition effects

While constraint differences usually don’t modulate N400 amplitudes in response to unexpected sentence continuations, differences in processing might be reflected in the post-N400 time window. The frontal positivity is observed starting at around 600 ms for plausible prediction violations in highly constraining sentences compared to merely unpredictable (but equally unexpected, i.e., matched cloze probability) words in weakly constraining sentences (Brothers et al., 2020; Federmeier et al., 2007; Kuperberg et al., 2020, but see also Stone et al., 2023) and is invariant to task demands (Lai et al., 2023). The pattern suggests a cognitive process dealing with the consequences of precise predictions that are violated. The exact functional specificity of the frontal positivity is still debated, but it generally seems to be related to additional effort necessary for representation update, by updating the underlying higher level message representation (Brothers et al., 2023; Kuperberg et al., 2020) or the suppression of the originally predicted sentence continuations (Ness & Meltzer-Asscher, 2018).The late frontal positivity is relevant to the discussion of adaptation processes in sentence repetition on two levels. As discussed above, parts of this experiment (i.e., the constraint manipulation) investigated a possible effect of constraint on changes in semantic predictions as reflected in the (reduced) N400 amplitude of repeated sentences. If constraint modulates N400 responses to repeated unexpected sentences, the mechanism driving this might be reflected in the initial frontal positivity. Additionally, the frontal positivity itself can be seen as an index of whether the consequences of unexpected information in a high constraint sentence change after processing the error. A potential mechanism could be changes in the higher-level expectations after processing the violation of a strong prediction, which reduces the cognitive control necessary to update message level representations downstream. Another possible explanation is that upon repetition, the most probable but previously incorrect continuation is less activated and less suppression is necessary (Ness & Meltzer-Asscher, 2018) or suppression is not required at all, as the frontal positivity is often described as all-or-nothing effect (e.g., Lai et al., 2023). A reduction of the frontal positivity when the same high constraint unexpected sentence is presented again later in the experiment would indicate less effort needed to deal with the unexpected continuation.

In repetition or memory tasks, the N400 is often followed by a late positive component with centroparietal distribution (LPC). The LPC is suggested to be related to more explicit memory processes (Rugg & Curran, 2007), deeper encoding (Paller et al., 1995; Paller & Kutas, 1992; Rugg et al., 1998) and recollection of episodic details (Wilding et al., 1995; Wilding & Rugg, 1996). Both, repeated sentences, and words repeated in new sentences, have previously been reported to show larger LPC amplitudes upon repetition (Besson et al., 1992; Besson & Kutas, 1993; Rugg, 1985; Rugg et al., 1998). This effect is stronger for repetitions of sentences with unexpected continuations (Besson et al., 1992) or repetitions of the unexpected word itself (Rommers & Federmeier, 2018b), compared to their expected counterparts. For words repeated in a new sentence, this enhancement doesn’t seem influenced by the initial sentence constraint (i.e., the precision of the prediction; Lai et al., 2021). Within this experiment we were interested in replicating LPC effects for unexpected item repetition across longer distances (compared to Rommers & Federmeier, 2018b) as well as in the exploration of the effect of constraint on the LPC in sentence repetitions.

### Time-frequency effects of repetition

Time frequency analysis of power allowed us to additionally replicate previous findings in literature using a sentence repetition paradigm and gain further insights regarding possible effects of constraint by capturing non-phase-locked signals, too. Repetitions of words within lists, usually combined with a recognition memory task, have previously been associated with theta (4-7 Hz) power increase and alpha (8-12 Hz) power decrease (Burgess & Gruzelier, 2000; Klimesch et al., 1997; Van Strien et al., 2007), whereas word repetition in new sentence contexts showed reduced power in the alpha and beta (13-30 Hz) frequency bands (Rommers & Federmeier, 2018b) upon repetition. The current experiment can add to these findings using repeated sentences, rather than word repetition only. Additionally, an investigation of possible repetition effects in the alpha/beta frequency band can supplement the analysis of the time-locked ERP signal in both the context and constraint manipulation.

Prediction violations by unexpected words or semantic anomalies in strongly constraining sentences often elicit an increase of power in the theta band, hypothesized to reflect control processes in response to error processing (Hald et al., 2006; Lewis & Bastiaansen, 2015; Pu et al., 2020; Rommers et al., 2017; Wang et al., 2012). A possible change of theta band power from the first presentation of high constraint sentences (violation of precise predictions) to their repetition can offer additional insights into the effect of repetition on the consequences of precise predictions that are violated. If theta power for highly constraining unexpected sentences is reduced at second compared to first presentation, this might indicate that changed expectations reduced or diminished the amount of cognitive control necessary to process the prediction violation. Additionally, research on predictive processing has been informed by pre-stimulus effects in the time-frequency domain. Alpha and/or beta band power was reduced prior to word onset in strongly constraining compared to weakly constraining sentences (Gastaldon et al., 2020; León-Cabrera et al., 2022; Li et al., 2017; Piai et al., 2014; Rommers et al., 2017; Wang et al., 2018; but see also: Rommers & Federmeier, 2018a; Terporten et al., 2019). These effects have been interpreted as a reflection of the predictive benefit of such sentences, with explanations for the underlying mechanism ranging from attentional preparation (Bidet-Caulet et al., 2012; Rommers et al., 2017) to lexical/memory access (Piai et al., 2014, 2020). Using time-frequency analysis, we therefore investigated the overall effect(s) of repetition and its interaction with context and constraint manipulations and tried to replicate pre-stimulus onset effects of predictive processing.

### Material and Methods

The study was pre-registered on the Open Science Framework (https://osf.io/r6kpf) and deviations are stated as such.

### Participants

The experiment is part of a research project whose protocols were approved by the Ethics Committee of the German Psychological Association (DGPs; MR102018). All participants gave written informed consent before the experimental session in accordance with the Code of Ethics of the World Medical Association (Declaration of Helsinki). 42 naive volunteers (9 male) participated in the experiment. The sample size was chosen to be 10 participants higher than other relevant and recently published literature (e.g., Lai et al., 2021; Rommers & Federmeier, 2018b) and pre-registered. Two participants had to be replaced due to excessive artefacts (< 25 trials in one or more conditions). Participants were compensated by either course credits or money (15€/h). Age of the final sample ranged from 18 to 36 years (M=23.09; SD=4.41). All were native speakers of German and right-handed (as assessed via the Edinburgh Handedness Inventory). All had normal or corrected-to-normal vision, and none reported a history of neurological or psychiatric disorders.

### Material and design

The study manipulated repetition (1st vs 2nd presentation) and constraint (high constraint vs. low constraint) in sentences with unexpected endings^1^. Within the low constraint unexpected sentences, repetition type was manipulated (sentence vs word in new sentence). Repetition type was only manipulated for low constraint sentences, as word repetition across different high constraint sentences would be (more) influenced by differences in semantic overlap between the strong prediction initially made and the unexpected continuation as well as contextual fit for the two sentence contexts chosen. In contrast, the comparison of low constraint sentences and the repetition of critical words in a new low constraint context allowed for a clean contrast that could be counterbalanced across participants (see table 1).

**Table 1.**
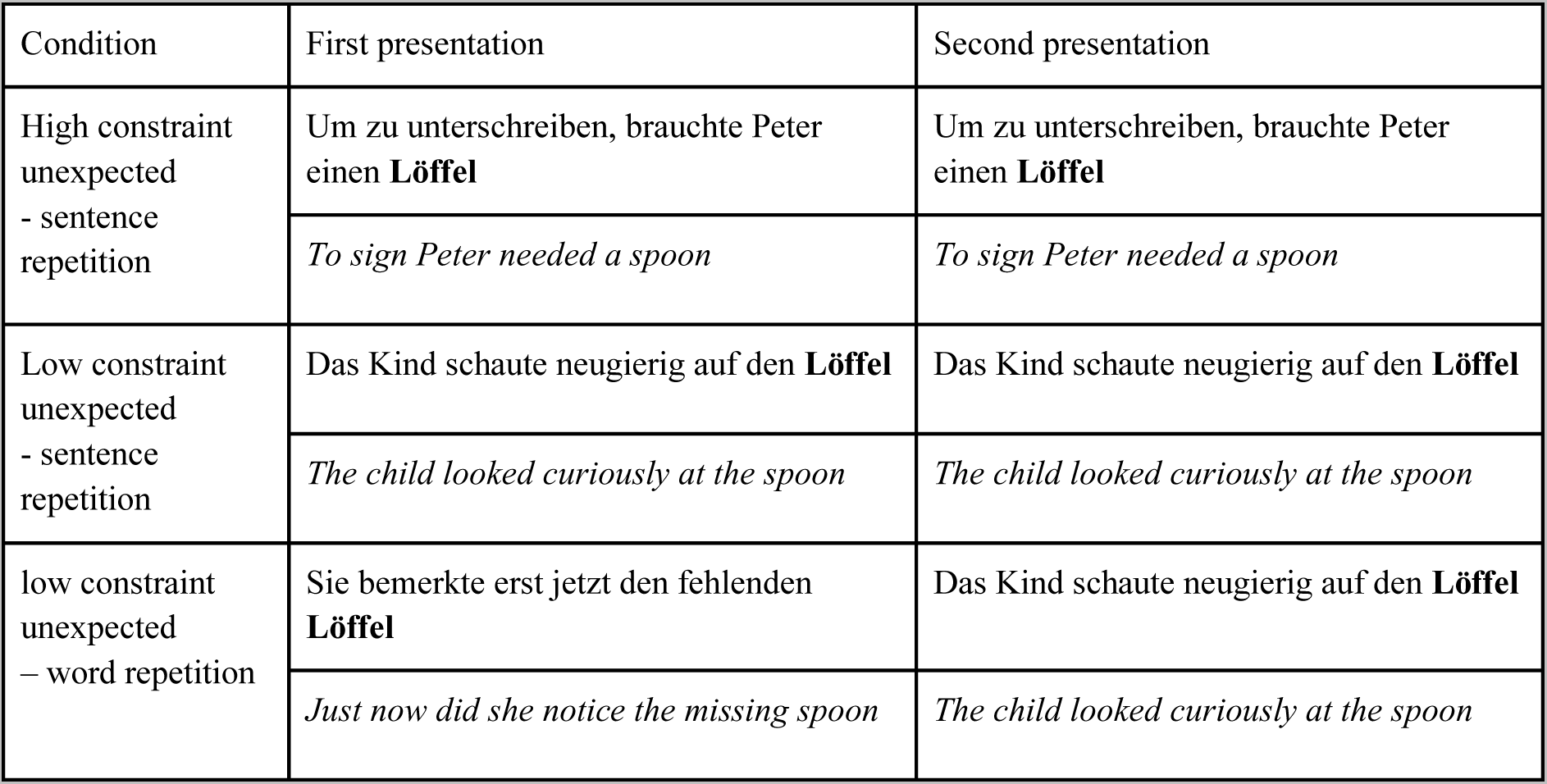
Conditions and example sentences with their approximate translation into English.

| Condition | First presentation | Second presentation |
| --- | --- | --- |
| High constraint unexpected<br>- sentence repetition | Um zu unterschreiben, brauchte Peter einen **Löffel** | Um zu unterschreiben, brauchte Peter einen **Löffel** |
|  | *To sign Peter needed a spoon* | *To sign Peter needed a spoon* |
| Low constraint unexpected<br>- sentence repetition | Das Kind schaute neugierig auf den **Löffel** | Das Kind schaute neugierig auf den **Löffel** |
|  | *The child looked curiously at the spoon* | *The child looked curiously at the spoon* |
| low constraint unexpected<br>– word repetition | Sie bemerkte erst jetzt den fehlenden **Löffel** | Das Kind schaute neugierig auf den **Löffel** |
|  | *Just now did she notice the missing spoon* | *The child looked curiously at the spoon* |

For two experiments comparing the repetition of previously unexpected words from high and low constraint sentences in a new low constraint sentence, and reporting no difference between these conditions, see Lai et al. (2021). Consequently, our stimuli consisted of 120 sentence triplets for the first presentation, two of which had low sentence constraint and one of which had strong constraint. All sentences of one triplet ended with the same unexpected word. For the strongly constraining sentences, the expected word was replaced with an unlikely but congruent continuation. The weakly constraining sentences all had congruent continuations, but did not end on the word with the highest cloze probability or a word semantically related to it (see Federmeier et al., 2007). The cloze probability of a word in a sentence context was derived from a sentence completion task, operationalized as the proportion of participants completing the sentence with a respective word. Sentence constraint was defined as the cloze probability of a sentence’s most frequently provided completion. The task was run online on Prolific (prolific.co) with 20 participants per list. Cloze probabilities for both, the high constraint and low constraint condition, were close to zero (high constraint: 0.01±0.03; low constraint: 0.04±0.07). Cloze probability of the most frequent completion (i.e., sentence constraint) was by definition higher in the high constraint (0.78±0.17) than in the low constraint sentences (0.26±0.10). Sentence length was matched (high constraint unexpected: 11.99 ± 2.67 words; low constraint unexpected: 11.64 ± 2.62 words). The sentences from each triplet were divided across three counterbalanced lists such that participants would see each unexpected word in only one of the conditions. 20 sentences later, the critical word was repeated. Depending on the condition, the critical word was repeated in the same sentence as previously (high constraint sentence repetition or low constraint sentence repetition) or in a new low constraint sentence context (low constraint item repetition). The second sentence in the low constraint item repetition condition corresponded to the other low constraint sentence from the target word triplet. The order and repetition type (sentence or item) for low constraint sentences was counterbalanced. For sentence examples see table 1.

In addition to that, 40 strongly constraining filler sentences with expected continuations were added to each list, 20 of which were repeated. These fillers were not analyzed but ensured an equal amount of highly constraining expected and unexpected sentences for each participant. The sentences in between repetitions comprised fillers as well as first presentations or repetition of sentences belonging to different triplets. The overall order of sentences (i.e., their first presentation) was randomized for each participant while additionally allowing only two high constraint unexpected sentences to be presented successively.

### Procedure

Participants were instructed to read silently for comprehension. Each trial started with a fixation cross. Participants started the sentence presentation via button press, which was followed by a 300 ms blank screen. Sentences were then presented word by word (190 ms + 20 ms for each letter) with a 300 ms interval between words (Rapid Serial Visual Presentation: RSVP). The sentence final word was presented for 500 ms for equal presentation times across critical words. After an additional interval of 800 ms the fixation cross appeared again. Stimulation was controlled by running the Psychophysics Toolbox (Brainard, 1997; Kleiner et al., 2007) in MATLAB R2020a (MathWorks Inc. Natick, MA, USA). Comprehension questions were asked following filler sentences only to ensure that participants were paying attention. For filler sentences that were repeated, a different question was asked upon the second presentation. The sentence presentation was divided into 8 blocks and participants were encouraged to take short breaks in between. Overall, the stimulus presentation lasted approximately 1h.

### Data acquisition and analysis EEG recording and analysis

While participants read the sentences their EEG was recorded at 32 active Ag/AgCl electrodes (actiCHamp, Brain Products) mounted in an elastic electrode cap based on the 10-20 system. Eye movements were monitored with bipolar electrodes at the outer canthi of the eyes. EEG and EOG were recorded with a sampling rate of 1000 Hz and electrode impedances were kept below 5 kΩ. Analyses were performed using EEGlab, ERPlab, Fieldtrip and MNE Python (Delorme & Makeig, 2004; Gramfort et al., 2013; Lopez-Calderon & Luck, 2014; Oostenveld et al., 2011). The EEG data was downsampled to 500 Hz and re-referenced to the average of the left and right mastoid. Channels with an accumulated z-score > 4 were removed and interpolated through spline interpolation (1 electrode for 17 of the subjects). The continuous data was then high pass filtered at 0.1 Hz (two-pass Butterworth with a 12 dB/oct roll-off). Eye movement, blinks, heart, and channel noise components (the last two not pre-registered but decided to be sensible upon inspection of the data) were corrected using independent components analysis (ICA) and the IClabel plugin (Pion-Tonachini et al., 2019). The IC weights were transferred from continuous data that was filtered at 1 Hz. IClabel then automatically removed all components that were classified as the respective category with a probability > 30 (average of 3.93 ±1.33 components per participant).

### ERP analysis

Data was low pass filtered at 30 Hz (two-pass Butterworth with a 24 dB/oct roll-off). The corrected signal was segmented from −200 to 1000 ms time-locked to stimulus onset.

Segments with values exceeding ±75 µV at the region of interest (ROI) channels were automatically removed. For data visualization all electrodes were evaluated for artefact rejection. On average 0.58±0.97 trials were rejected per participant. As recommended by Alday (2019) the 200 ms baseline period was included into our statistical model. For the statistical analysis mean amplitudes were extracted in an a priori determined time window of 300-500 ms after stimulus onset, averaged across a centro-parietal ROI where N400 amplitude is usually maximal (Cz, CP1, Cp2, P3, Pz, P4; see e.g., Kuperberg et al., 2020; Rommers & Federmeier, 2018a). Additionally, mean amplitudes were extracted in the 600-800 ms time window at the centro-parietal ROI, as typical for the LPC (e.g., Lai et al., 2021; Rommers & Federmeier, 2018b) and in the same 600-800 ms time window but in a frontal ROI (F3, Fz, F4, FC1, FCz, FC2; e.g., Kuperberg et al., 2020) to investigate the frontal positivity (not pre-registered).

The single trial data was analyzed using Bayesian linear mixed effects models using the ‘brms’ package (Bürkner, 2017) which fits Bayesian multilevel models in the Stan programming language (Stan Development Team, 2018). We accounted for the possibility of slight differences at the first presentation, as well as the inevitable difference in sentence stimuli between conditions for the constraint manipulation, by operationalizing each of our hypotheses in terms of repetition effect (first vs. second presentation) instead of analyzing ERP amplitudes at second presentation only. We first analyzed the effect of repetition type (sentence vs word in new sentence) and repetition, and then the effect of constraint (high vs low constraint) and sentence repetition, as the repetition type manipulation was nested within the low constraint condition. The effect of interest for each hypothesis is the respective interaction effect. We fit the maximal model with by-subject and by-item random slopes for all fixed effects of interest as well as their interaction. All categorical fixed effects were sum coded (first: -0.5, second: 0.5/high constraint: -0.5, low constraint: 0.5/item repetition: -0.5, sentence repetition: 0.5). The baseline interval was added as an additional fixed effect (Alday, 2019), as well as the cloze probability as a nuisance factor. We included normally distributed, regularizing priors for the intercept (mean = 0, SD = 5) and for all fixed effects as well as their interactions (mean = 0, SD = 1). We also defined priors for subject and item variability (mean = 0, SD = 0.5), residual variance (mean = 8, SD = 2) and correlations of random effects using a LKJ(2) prior (Bürkner, 2017; Lewandowski et al., 2009). These priors were informed by a previous Bayesian ERP analysis, which suggests higher intercept variability than individual variability between participants and items (Nicenboim et al., 2020) and ensured stable and physiologically plausible estimates (Chung et al., 2015; Gelman et al., 2008, 2017). Models were fit with 20,000 iterations (incl. 2000 iteration warmup; the number of iterations had to be increased compared to the pre-registration to obtain stable Bayes factors) and model convergence was evaluated by making sure that the number of bulk and tail effective samples for every parameter estimate was at 10% of the post-warmup samples (=1800 samples) and that R-hat values did not exceed 1.01. Support for or against the outlined hypotheses was assessed using Bayes factors. These Bayes factors were calculated using bridge sampling (Bennett, 1976; Gronau et al., 2017; Meng & Wong, 1996). We compared models with the predictor of interest versus a reduced model without this predictor (i.e., BF10) and evaluated the strength of evidence in reference to Jeffrey’s scale (1939).

Using this convention, as the Bayes factor increases over 3, evidence strengthens against the null hypothesis. As the Bayes factor decreases under 0.3, evidence strengthens in favor of the null hypothesis. Given the sensitivity of the Bayes factor to the choice of prior (Lee & Wagenmakers, 2014), Bayes factors were calculated for a range of different priors on the effects of interest, while holding all other priors constant. The priors for the sensitivity analysis ranged from Normal(0,.2) to Normal(0,3).

### Time-frequency analysis

Epochs were extracted from -1700 to 1200 ms relative to the onset of the final word of each sentence (incl. 200 ms data padding). Artefacts were rejected following the same steps as listed for the ERP analysis. Separate analyses were conducted of the signal around the onset of the critical word (−500 to 1000 ms) and the signal preceding the critical word (−1500 to 0 ms) (see Rommers et al., 2017 for a similar approach). Line noise was removed using cleanLineNoise from the PREP-Pipeline (Bigdely-Shamlo et al., 2015) and data was low pass filtered at 100 Hz. Time frequency representations were derived using complex Fourier coefficients for a moving time window (fixed length of 0.01 s; Hanning taper; moving in steps of 0.4 s) for frequencies 2-30 Hz with a resolution of 1 Hz. The resulting spectrograms were averaged within participant and condition. For contrasts between the conditions, each condition was divided (elementwise) by the average power spectrogram across all conditions, to prevent differences in pre-stimulus baseline activity (due to differences in constraint) from influencing power estimates later in the epoch and allow for investigation of pre-stimulus activity (León-Cabrera et al., 2022; Piai et al., 2014; Rommers et al., 2017). To visualize the power changes for the individual conditions, we computed power changes relative to the baseline from -500 to -150 ms time-locked to the onset of the stimulus. Given the conflicting results in existing literature concerning time points, channels, and frequency bands in which the reported effects occur, statistically significant differences between conditions were identified using nonparametric cluster-based permutation tests (Maris & Oostenveld, 2007) and are reported as an exploratory analysis. In the cluster-based permutation test, data triplets (electrode x time x frequency) were compared between two conditions based on a paired-sample t-statistics. Channels had 7.9 neighbors on average and neighboring triplets which exceeded the critical α-level of 0.05 formed a cluster. The permutation p-value was computed using the Monte Carlo method involving 1000 random permutations of the same dataset. Permutation p-values were doubled for a two-tailed test, which allowed us to evaluate the significance of clusters at α = 0.05.

## Results

On average participants answered 96 ± 0.013% of the questions correctly, which indicates that they were paying attention throughout the experiment.

### ERP analysis: Manipulation of sentence context and repetition

Figure 2 plots the grand average of the first presentation of low constraint sentences compared to a repetition of the same sentence or a new low constraint sentence with the repeated final word to explore the effect of context on N400 repetition effects. Probability densities and corresponding Bayes factors are visualized in Figure 4A. The Bayes factor shows extreme evidence for an effect of repetition (*β* = 0.65 [0.34, 0.96], BF10 = 463), but also moderate evidence for an interaction effect between repetition and context (*β* = 0.65 [0.06, 1.24], BF10 = 3.16). The sensitivity analysis for the interaction effect can be seen in Figure 4C. This interaction is driven by the more positive voltage in the repeated sentence condition as compared to the repetition of a word in a new sentence, as visible in the grand average plot (Figure 2A). When a sentence is repeated, the N400 is reduced (more positive) compared to the first presentation of the sentence (M = 0.973 [0.54, 1.40]; pairwise post-hoc comparison: BF10 = 1230). There was inconclusive evidence regarding a repetition effect when only analyzing words that were repeated with a new context (M = 0.33 [-0.10, 0.76]; pairwise post-hoc comparison: BF10 = 0.60). At the first presentation the conditions didn’t differ, so we would not expect a main effect of repetition type (*β* = 0.34 [0.01, 0.66], BF10 = 1.3).

**Figure 2.**
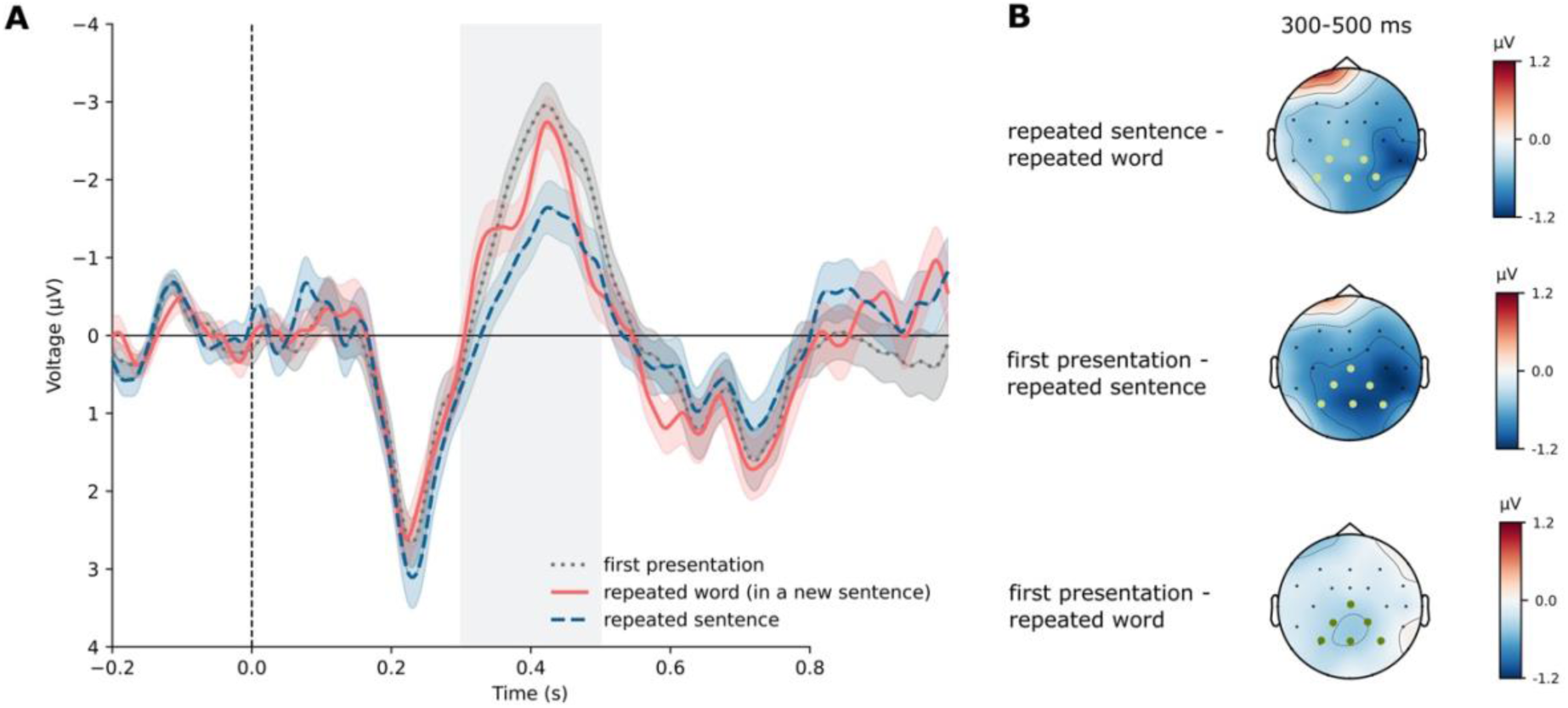
(A) Grand-average waveforms (n = 42) at centro-parietal ROI with regression-based baseline correction based on the 200 ms before stimulus onset. The 300 to 500 ms time window is shaded in gray. Error bands indicate the SEM. (B) Topographies for the 300 to 500 ms time window. The centro-parietal ROI is marked with green dots.

In the post N400 time window, data was analyzed with respect to a possible LPC in a centro-parietal ROI, usually associated with explicit memory upon repetition. However, as can be seen in figure 2, there seem to be no condition differences in the late time window of interest. This was supported by the Bayesian analysis. There was moderate evidence against an effect of repetition (*β* = 0.09 [-0.22, 0.41], BF10 = 0.19), as well as against an interaction of repetition and repetition type, i.e., item versus sentence repetition (*β* = -0.09 [-0.69, 0.50], BF10 = 0.31) on the LPC. We would not expect a main effect of repetition type since sentences didn’t differ at their first presentation (*β* = -0.22 [-0.55, 0.10], BF10 = 0.44).

### ERP analysis: Manipulation of sentence constraint and repetition

Figure 3A illustrates that repeated unexpected sentences elicit a smaller (more positive) N400 for both low constraint and high constraint sentences, replicating the classic N400 repetition effect.

**Figure 3.**
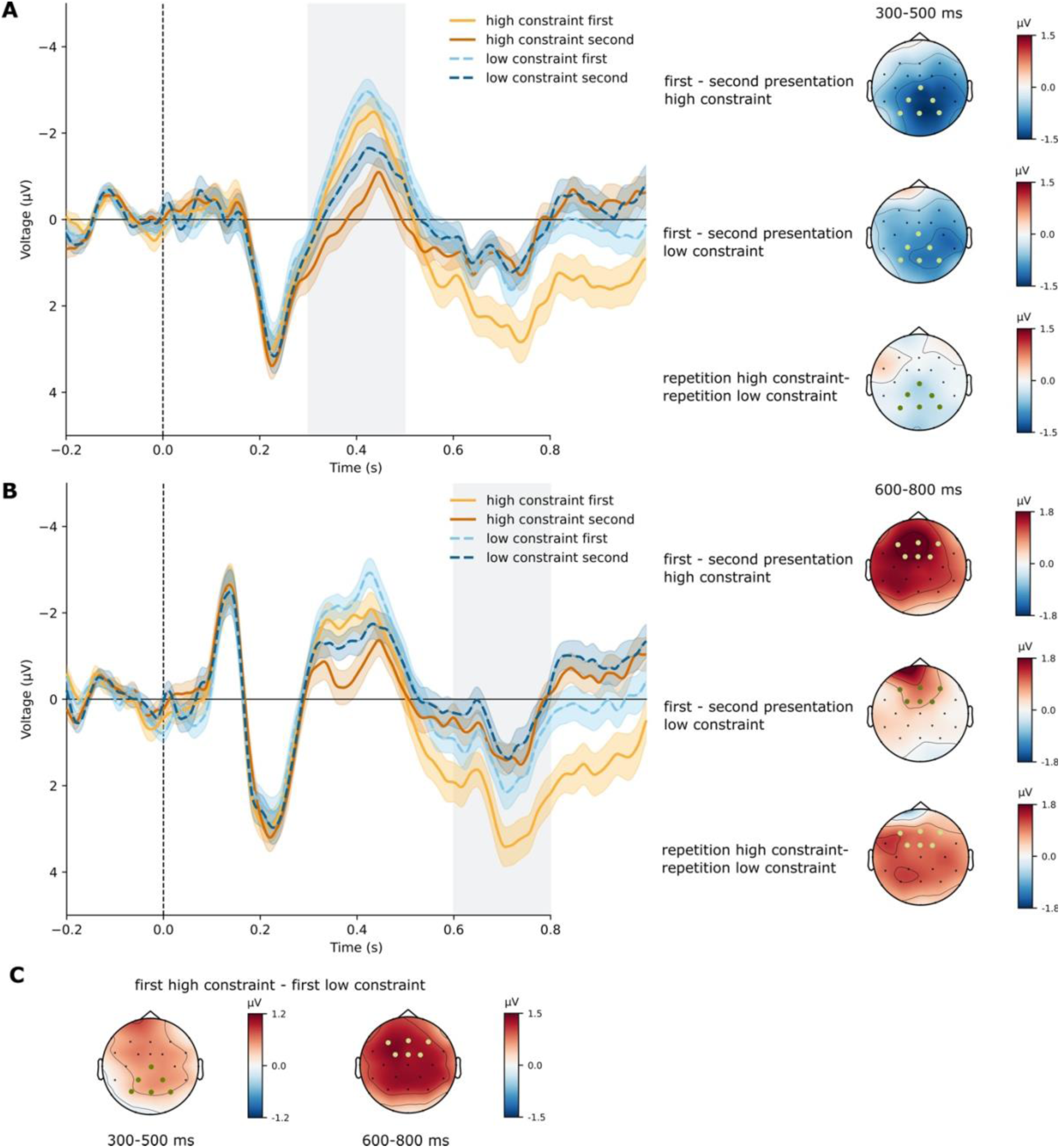
(A) Grand-average waveforms (n = 42) at centro-parietal ROI with regression-based baseline correction based on the 200 ms before stimulus onset. The 300-500 ms time window is shaded in gray. Error bands indicate the SEM. Topographies for the 300 to 500 ms time window show the repetition effects and their interaction. The centro-parietal ROI is marked with green dots. (B) Grand-average waveforms at frontal ROI with regression-based baseline correction based on the 200 ms before stimulus onset. The 600-800 time window is shaded in gray. Error bands indicate the SEM. Topographies for the 600 to 800 ms time window show the repetition effects and their interaction. The frontal ROI is marked with green dots (C) Topographies for the comparison between the first presentation of high constraint sentences and low constraint sentences in the time window corresponding to the N400 (300-500 ms) and to the frontal positivity (600-800 ms), respectively.

**Figure 4.**
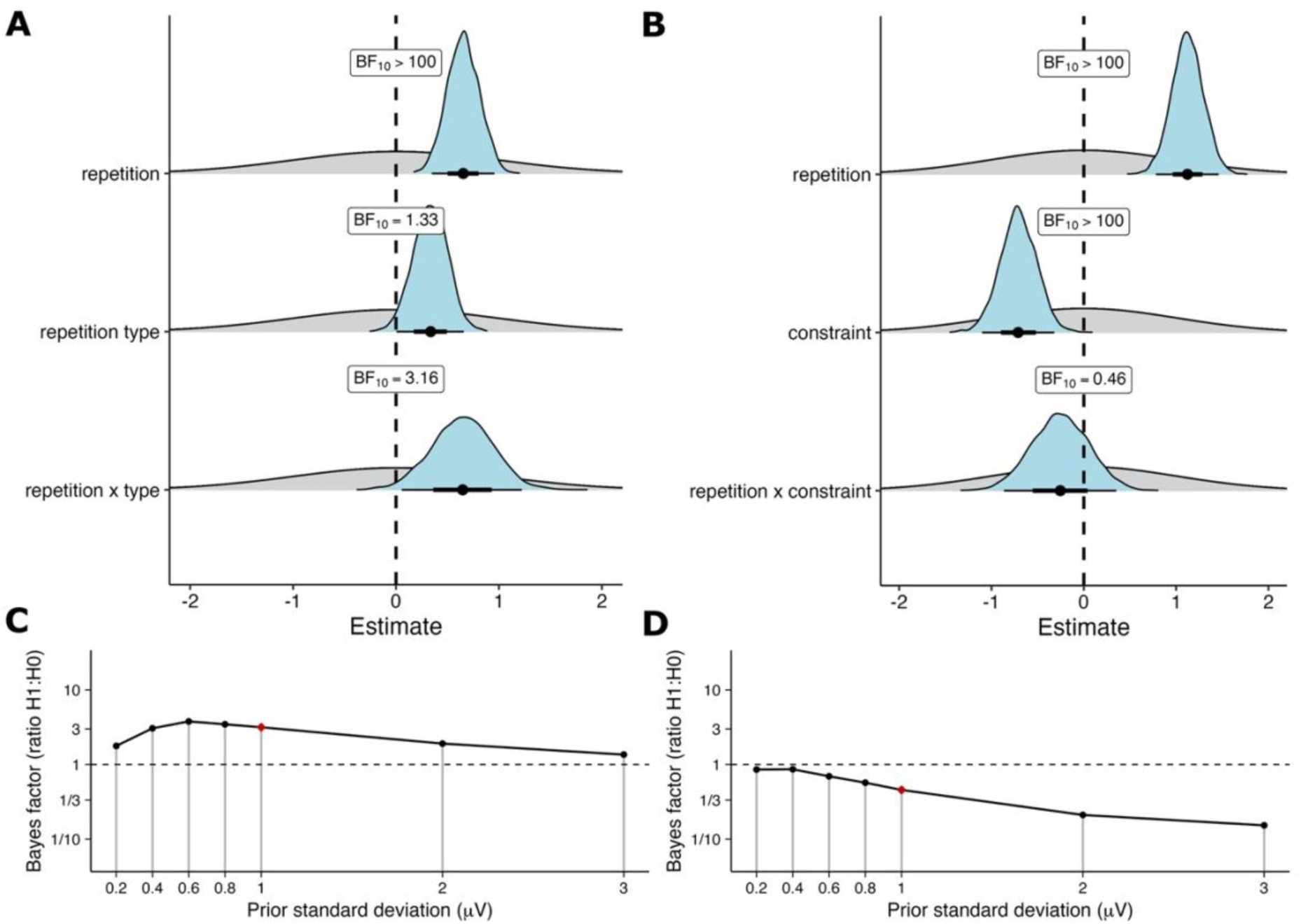
(A) The effect of repetition type (sentence or word in new context) on repetition: Posterior (blue) plotted with prior density (gray). (B) The effect of constraint (high or low) on sentence repetition: Posterior (blue) plotted with prior density (gray). (C) Sensitivity analysis for the interaction of repetition and repetition type (sentence or word in new context). (D) Sensitivity analysis for the interaction of repetition and constraint (prediction violation). (C+D) The horizontal line at a Bayes factor of 1 in each plot indicates equal evidence for H_1_ and H_0_. Above this line, evidence increases for H_1_ (evidence for an interaction effect), below this line, for H_0_ (evidence against an interaction effect).

Results from the Bayesian mixed-effect model are reported in Figure 4B. The Bayes factors show extreme evidence for an effect of repetition (*β* = 1.12 [0.78, 1.46], BF10 = 2006057). Repeated sentences elicited a reduced N400 compared to the first presentation in both highly constraining (second minus first presentation: M = 1.25 [0.79, 1.70]) and weakly constraining sentences (M = 1.0 [0.54, 1.44]). Unexpectedly, there was also extreme evidence for an effect of constraint (*β* = -0.70 [-1.09, -0.32], BF10 = 109). Specifically, low constraint sentences elicited a more negative N400 both at their first (low constraint minus high constraint: M = - 0.58, [-1.07, -0.10]) and the second presentation (M = -0.84 [-1.32, -0.35]). There was anecdotal evidence against an interaction of repetition and constraint with a BF10 of 0.46 (*β* = -0.26 [-0.85, 0.33]). This result reflects that there is 1/0.46 = 2.17 times more evidence for the null model without the interaction as predictor. The sensitivity analysis for the interaction effect can be seen in figure 4D.

In the post-N400 time window there were two ERP components of interest: the frontal positivity as a consequence of precise violated predictions and the LPC in a centro-parietal ROI usually associated with stimulus repetition.

#### Frontal positivity

As can be seen in figure 3B, the first presentation of highly constraining unexpected sentences (violation of precise predictions) seems characterized by a late frontal positivity. Statistically, the analysis within the frontal ROI led to moderate evidence for an interaction between repetition and constraint (*β* = 0.85 [0.16, 1.54], BF10 = 6.04). There was also strong evidence for a main effect of constraint (*β* = -0.61 [-0.99, -0.23], BF10 = 20.59) and repetition (*β* = -1.15 [-1.59, -0.70], BF10 = 15325). At their first presentation, high constraint unexpected sentences were more positive than low constraint unexpected sentences (M = 1.04 [0.52, 1.55]; pairwise post-hoc comparison: BF10 = 367). This difference was reduced upon sentence repetition (M = 0.19 [-0.33, 0.70]; pairwise post-hoc comparison: BF10 = 0.30).

#### LPC

Repeated sentences were not characterized by a visually more positive going ERP in the post N400 time window in the centro-parietal cluster, as would have been expected for an LPC effect (Fig. 3). Regarding the effect of constraint, figure 3A illustrates that differences between the conditions are mainly driven by the frontal positivity in the first presentation of high constraint unexpected sentences, which is partly also reflected in the centro-parietal electrodes. The results from the centro-parietal ROI therefore merely mirror those from the frontal positivity above. This leads to strong evidence for an interaction of repetition and repetition type, i.e., high constraint versus low constraint sentence repetition (*β* = -0.91 [- 0.29, 1.54], BF10 = 18.83). Highly constraining unexpected sentences were more positive than weakly constraining unexpected sentences at first presentation (M = 1.12 [0.7, 1.62]).

This difference was reduced upon sentence repetition (M = 0.24 [-0.22, 0.70]). Overall, there was inconclusive evidence regarding an effect of repetition (*β* = -0.44 [-0.88, 0.01], BF10 = 1.26) and extreme evidence for an effect of constraint (*β* = -0.69 [-1.03, -0.36], BF10 = 406.66). Even though these effects were (also) obtained in the centro-parietal ROI, it is important to note that they do not reflect LPC effects per se, but rather originate from the frontal positivity.

### Exploratory analysis: component overlap

Due to the unexpected main effect of constraint (see Fig. 1B for a visualization of the expected pattern), we examined the topography of high versus low constraint sentences at their first presentation in both the N400 and the late time window (600-800 ms) corresponding to the post N400 frontal positivity (Brothers et al., 2020; Federmeier et al., 2007; Kuperberg et al., 2020). The topographies reveal a striking similarity in that the maximum is left frontal in both time windows (rather than centro-parietal, as would be typical for the N400; Fig 3C). The unexpected difference in N400 amplitude between the constraint conditions might thus stem from a spatio-temporal component overlap: the increased frontal post-N400 positivity for high constraint unexpected sentences as compared to low constraint unexpected sentences in our data could start earlier for some participants or trials resulting in more positivity and thus an attenuated N400 for highly constraining sentences (see also Brouwer & Crocker, 2017 and Delogu et al., 2021 for similar considerations regarding the N400 and P600). To account for this possibility, we conducted an exploratory temporal decomposition of the single trial ERP using the RIDE (Residue Iteration Decomposition) toolbox (Ouyang et al., 2011). RIDE builds on the assumption that ERPs can be disentangled into different components of either consistent or variable delays (Ouyang et al., 2015). This allows to control and correct for the variable latency of components across trials and participants to better isolate different components despite this latency variability. It has previously been shown that RIDE can separate component clusters associated with stimulus and response (Ouyang et al., 2013; Stürmer et al., 2013; Verleger et al., 2014) but also overlapping ERPs such as N400 and P600 (Ouyang et al., 2016). The RIDE algorithm requires baselined single trial data per subject and condition (Ouyang et al., 2015). Data was therefore baseline corrected from -200 to stimulus onset. Besides the stimulus related time window (S: 0-400 ms), two component clusters were included with time windows corresponding to the N400 (C1: 200-600) and the late frontal positivity (C2: 400-1000). These time windows must be larger than their traditional ERP time window to account for temporal variability of the single trial data (see e.g., Mauchand et al., 2021 for similar time windows). After extraction, components were re-baselined (Ouyang et al., 2015). Single trial data was reconstructed from the estimated components (following the protocol by Takács et al., 2022) and mean amplitudes for statistical analysis were calculated analogous to non-decomposed data as described above. A grand average plot of the two estimated components can be seen in figure 5. The algorithm successfully extracted both the N400 and the late frontal positivity.

**Figure 5.**
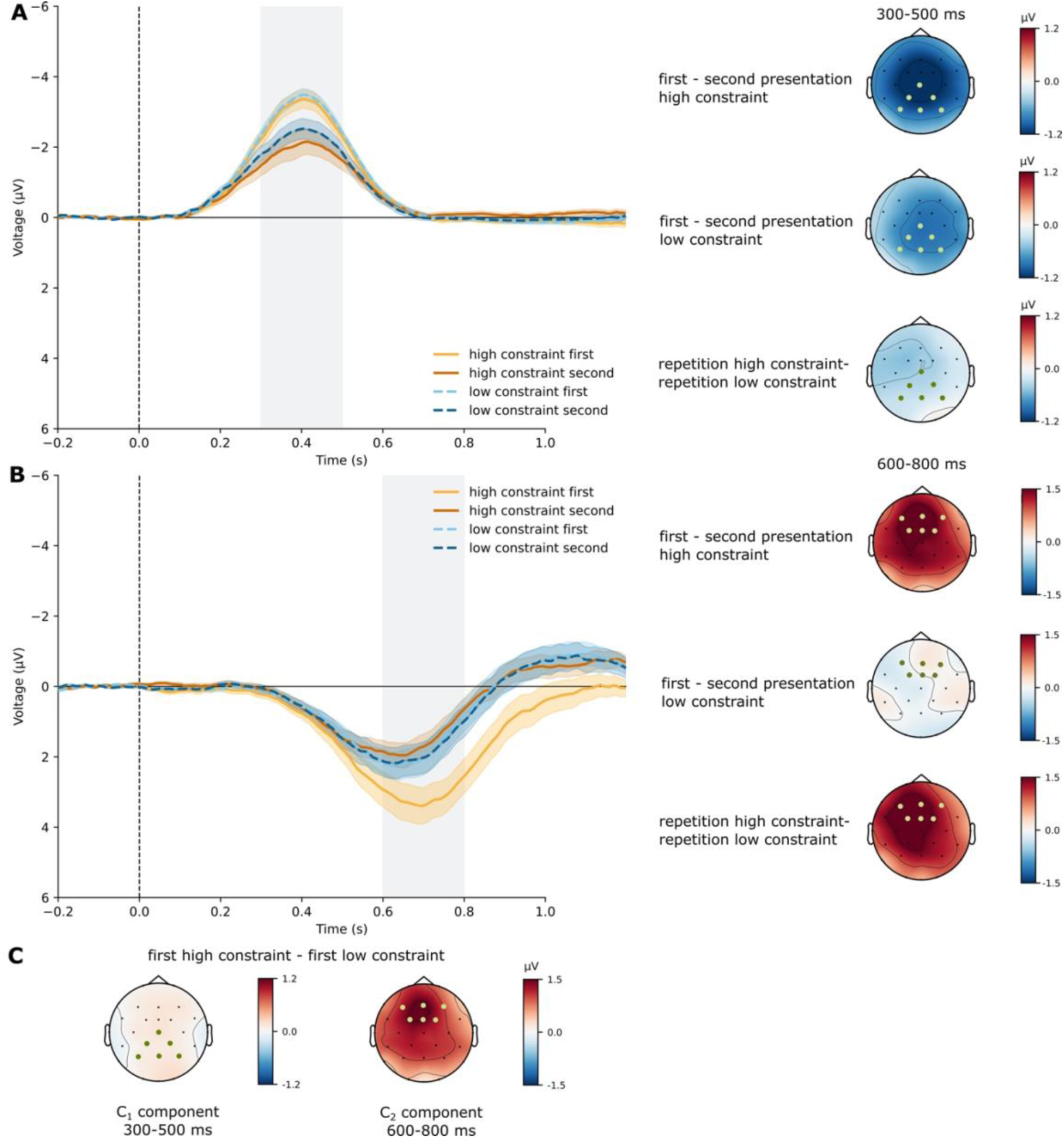
(A) Grand-average waveforms (n = 42) of RIDE component C_1_ at centro-parietal ROI. The 300 to 500 ms time window is shaded in gray. Error bands indicate the SEM. Green dots in topographies indicate centro-parietal ROI. (B) Grand-average waveforms (n = 42) of RIDE component C_2_ at frontal ROI. Error bands indicate the SEM. Green dots in topographies indicate frontal ROI. (C) Topographies for the comparison between the first presentation of high constraint sentences and low constraint sentences corresponding to theC_1_ component between 300 and 500 ms and to the C_2_ component between 600 and 800 ms, respectively.

Using this isolated N400 component, Bayes factor analysis again showed extreme evidence for an effect of repetition (*β* = 0.95 [0.46, 1.4], BF10 = 208). Repetition attenuated the N400 for highly constraining (M = 1.05 [0.44 1.66]) and weakly constraining sentences (M = 0.85 [0.26 1.44]). Using the RIDE component, there was inconclusive evidence for an influence of constraint (M = -0.43 [-0.88, 0.02], BF10 = 1.30). Low constraint sentences still elicited a numerically more negative amplitude in the first (M = -0.33 [-0.92 -0.22]) and the second presentation (M = -0.53 [-1.11 0.04]), but the difference was reduced compared to the raw ERP. There was anecdotal evidence against an interaction of repetition and constraint with a BF10 of 0.42 (*β* = -0.19 [-0.91, 0.51]), indicating 2.38 times more evidence for the null model without the interaction as predictor. Bayesian mixed-effect model results are illustrated in figure 6.

**Figure 6.**
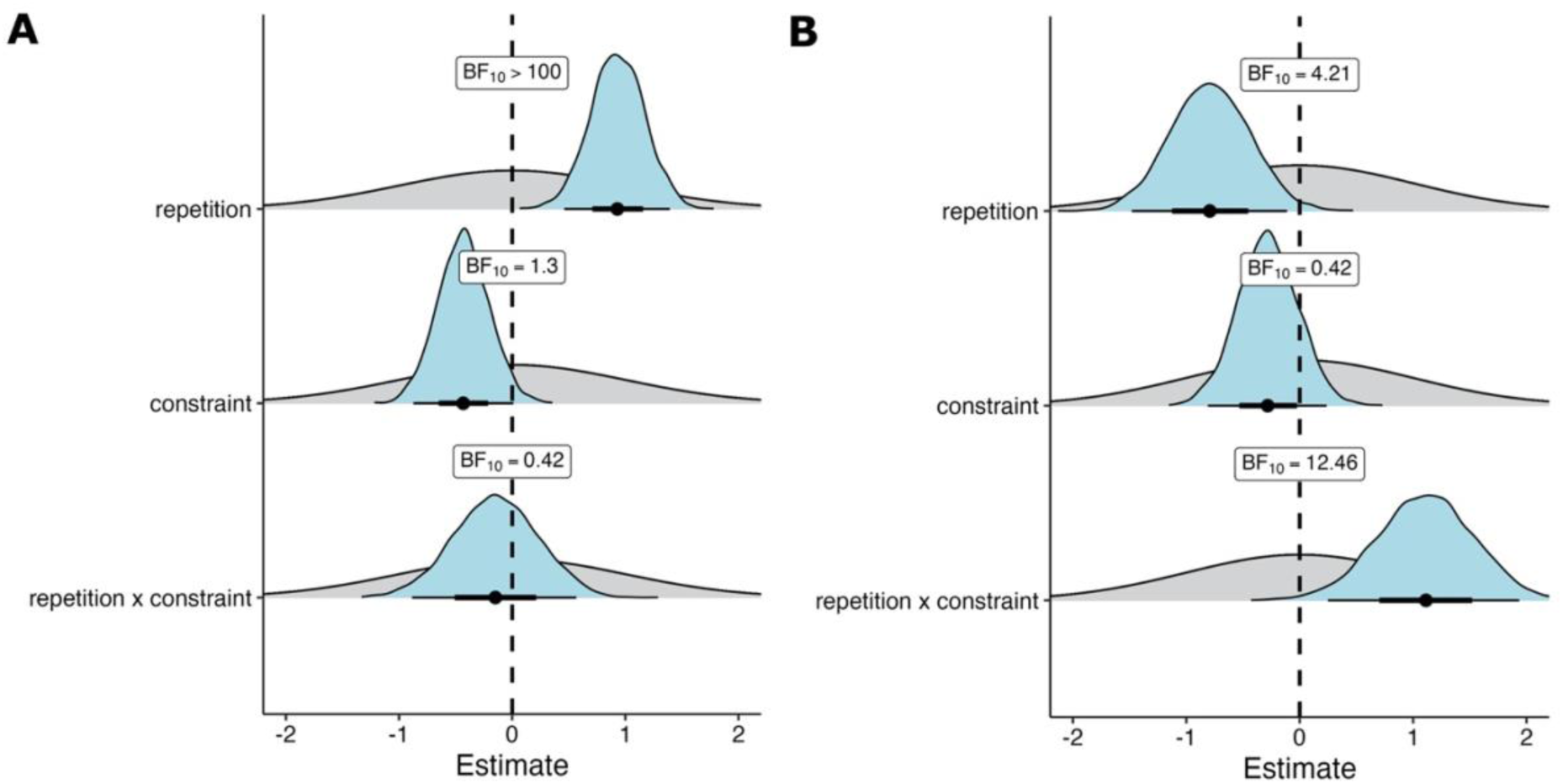
The effect of constraint on sentence repetition for RIDE components: Posterior (blue) plotted with prior density (gray). (A) C_1_ component (N400) (B) C_2_ component (late frontal positivity).

An analysis of the isolated frontal positivity supported the results from the raw ERP data. There was strong evidence for an interaction between repetition and constraint (*β* = 1.11 [0.27 1.95], BF10 = 12.46). There was anecdotal evidence against a main effect of constraint (*β* = - 0.28 [-0.80, -0.24], BF10 = 0.42) and moderate evidence for an effect of repetition (*β* = -0.80 [-1.44 -0.11], BF10 = 4.21). At their first presentation, highly constraining unexpected sentences were more positive than weakly constraining unexpected sentences (M = 0.84 [0.17 1.49]; pairwise post-hoc comparison: BF10 = 6.68). However, this effect was diminished upon repetition (M = 0.28 [-0.39 0.95]; pairwise post-hoc comparison: BF10 = 0.43).

### Time-frequency analysis: Manipulation of sentence context and repetition

A time-frequency analysis of the context manipulation revealed no significant differences between the first presentation of an unexpected word in a low constraint sentence and the repetition of this unexpected word in a new low constraint sentence (all clusters *p*>0.8; Fig. 7A). In contrast, the comparison of low constraint sentences at their first presentation and repetition of the complete sentence (Fig. 7B) indicated an effect spread throughout the post-stimulus time window, but clusters were most prominent between 400-1000 ms^2^, where power in the alpha and beta frequencies (8-22 Hz) was decreased upon second presentation. This effect exhibited a central distribution, lateralized to the right in the post 400 ms time window. Two significant positive clusters were found (*p*=0.02 and *p*=0.002). The contrast between the two repetition effects (repetition of sentence – repetition of word in new sentence context) revealed a significant interaction effect (one significant cluster *p*=0.03). Repeated sentences were characterized by a stronger reduction of power in the alpha and beta band (8- 22 Hz), most prominent after 600 ms and with a central distribution.

**Figure 7.**
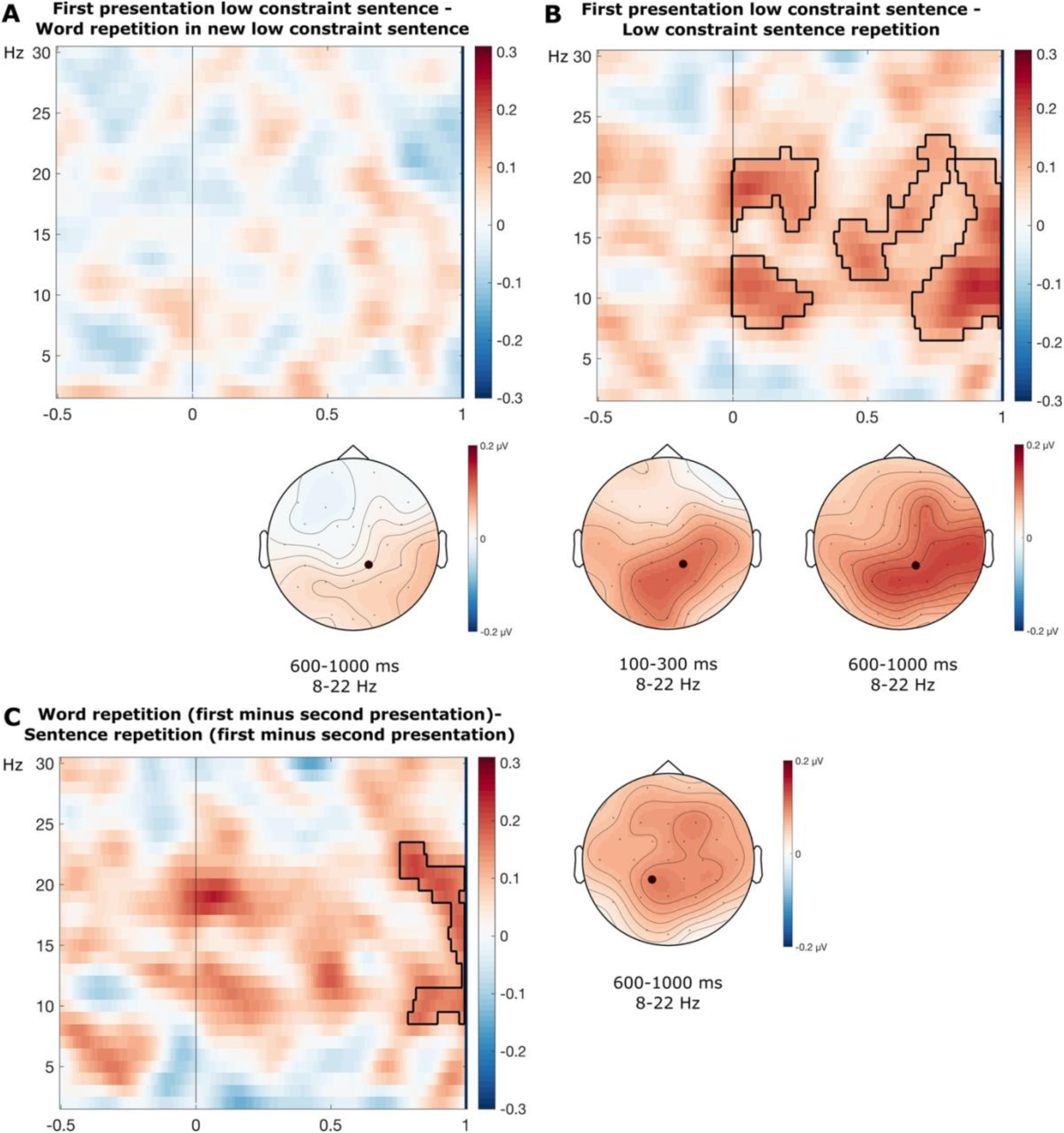
Grand average time-frequency plots of power changes. Time zero indicates the onset of the target word. The contour line in the spectrogram indicates the extend of clusters in a permutation test of the difference. (A) Contrast between first presentation of an unexpected word and its repetition in a new low constraint sentence at CP2, indicated with a black dot in the topography. (B) Contrast between the first presentation of an unexpected word in a low constraint sentence and its repetition in the same sentence context (sentence repetition) at CP2. (C) Interaction between repetition and repetition type (sentence vs word in new sentence) at CP1, indicated with a black dot in the topography.

### Time-frequency analysis: Manipulation of sentence constraint and repetition

Power changes at initial presentation of the high and low constraint sentences with unexpected endings can be seen in figure 8A. Sentence endings for both constraint conditions elicited an early broadband power increase with an occipital maximum, followed by a central (lower) theta power increase and an alpha/beta decrease with a broad centro-parietal distribution. As hypothesized, strongly constraining sentences with an unexpected continuation seem to exhibit a stronger theta power increase with a fronto-central distribution than weakly constraining sentences at first presentation (Fig. 8B). A cluster in the observed data was found around 400 to 800 ms, most consistently spanning frequencies between 3 and 7 Hz. A cluster-based permutation test indicated that there was a significant difference between the two conditions at first presentation (one significant positive cluster *p*=0.002).

**Figure 8.**
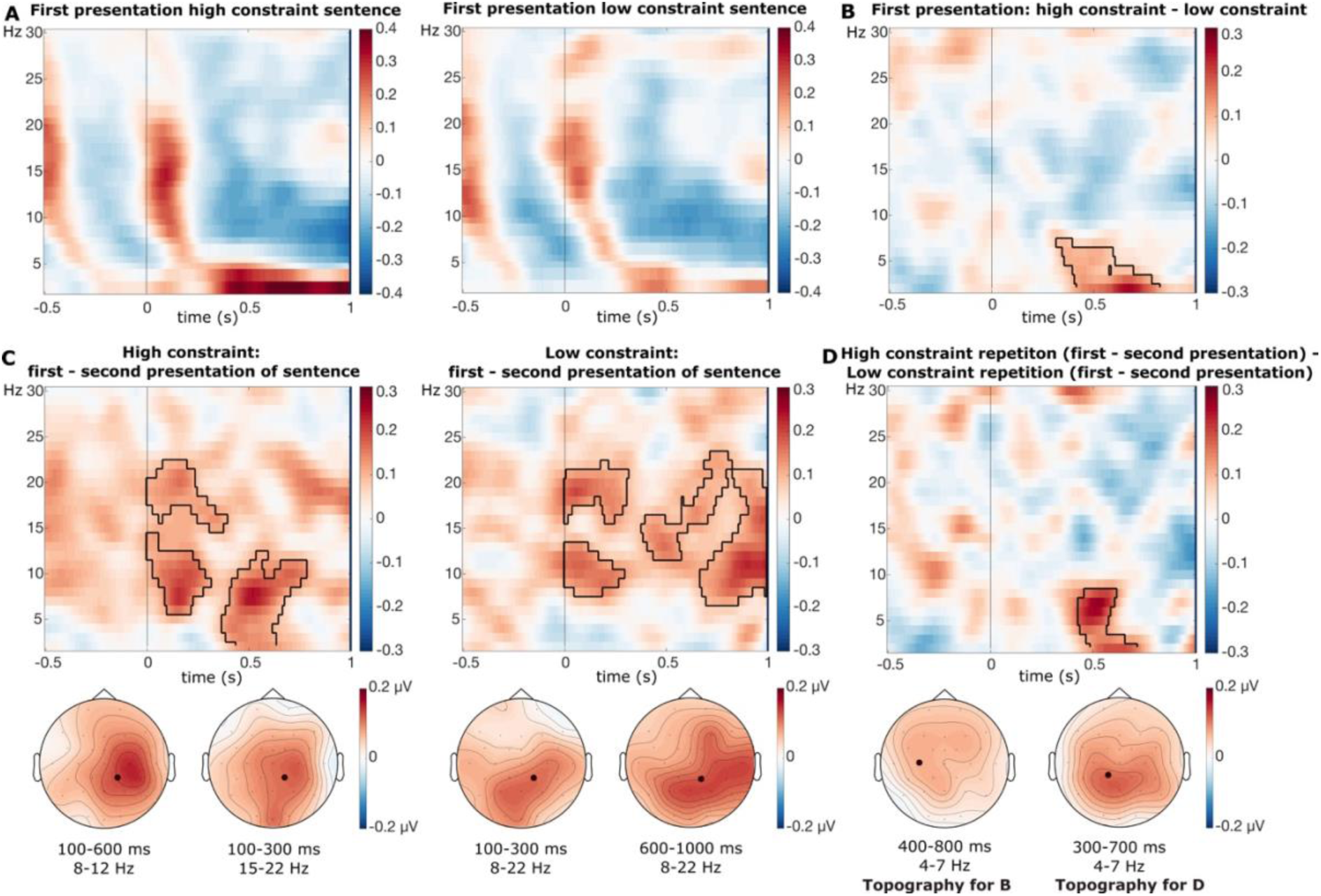
Grand average time-frequency plots of power changes. Time zero indicates the onset of the target word. The contour line in the spectrogram indicates the extend of clusters in a permutation test of the difference. (A) Individual conditions at CP2. The color scale indicates the proportion power change relative to a −500 to −150 ms baseline. (B) The effect of constraint at the first sentence presentation at C3, indicated with a black dot in the topography. (C) Contrast between the first and the second presentation of the sentences at CP2, indicated with a black dot in the topography. Note that the low constraint sentence repetition is the same figure as 7B but is repeated for ease of comparison. (D) Interaction between repetition and constraint at CP1, indicated with a black dot in the topography.

Figure 8C shows the effect of sentence repetition in the time-frequency domain at the onset of the sentence final word for high and low constraint sentence repetitions separately. For strongly constraining sentences, the comparison (first presentation minus second presentation) indicated an early power difference in the alpha/beta band, followed by an effect in the theta and alpha frequencies. Power in these frequencies was lower at the second presentation of the sentence. This effect exhibited a central distribution, lateralized to the right. More specifically, clusters spanned activity across the epoch and most consistently including frequencies between 2 and 20 Hz. Two significant positive clusters were found for this comparison (*p*=0.01 and *p*=0.006). The results for the same comparison for low constraint sentences have been reported above with respect to the context manipulation (see also Fig 7B). An analysis of potential interaction effects between constraint and repetition (first minus second presentation high constraint versus first minus second presentation low constraint; Fig 8D), also revealed a difference in the theta band. This effect can be explained by the pronounced theta band effect at the first presentation of the highly constraining sentences (Fig 8B) that was reduced upon repetition. The difference was most prominent around 400 to 700 ms and frequencies between 2 and 7 Hz. One significant positive cluster was found (*p*=0.03). Since the theta band effect seemed to mirror the pattern (with respect to both conditions and timing) exhibited by the frontal positivity (Fig. 3A) in the ERP, an additional analysis explored the extent to which the observed differences might be driven by phase-locked activity. For this, the ERP signal (phase-locked activity) was subtracted from the data of each participant and condition (Cohen, 2014), leaving only non-phase locked data. The theta band synchronization for prediction violations in high constraint sentences compared to low constraint unexpected words at first presentation was supported by a significant cluster with a similar distribution in non-phase-locked data (*p*=0.002). For the comparison of repetition effects, the effect in the theta band did not reach significance (all clusters *p*>0.4).

Interestingly, no significant difference (all clusters *p*>0.7) between highly and weakly constraining sentences at their first presentation was found in the alpha or beta frequency band in the pre-stimulus time window (Fig. 9).

**Figure 9.**
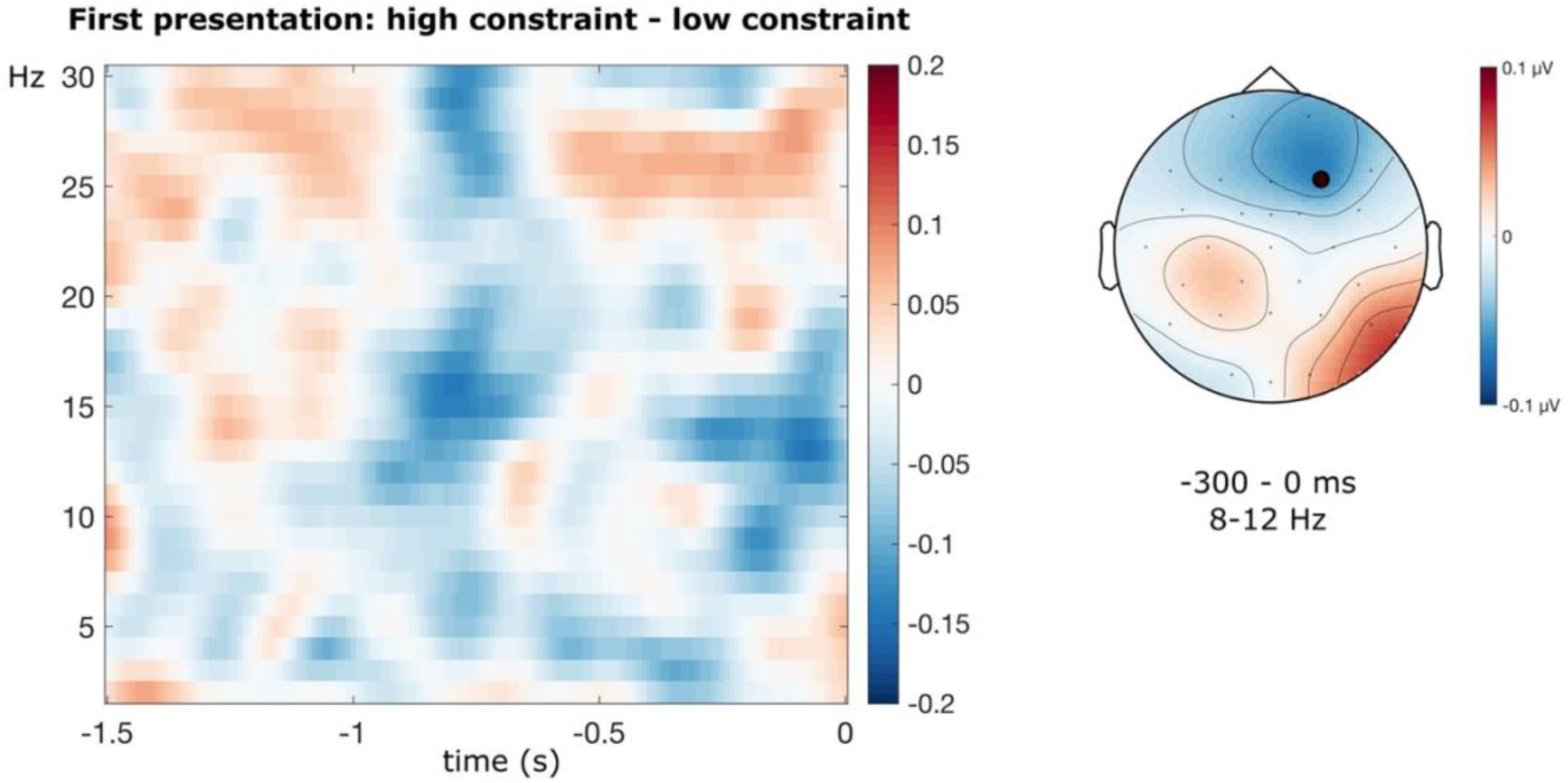
Comparison of pre-stimulus power changes. Grand average time-frequency plots of power changes at F4, indicated with a black dot in the topography.

## Discussion

The current study investigated how predictions in language processing adapt after processing unexpected information. Specifically, we examined whether predictions made from sentence context are updated when a continuation was unexpected, beyond changes to the representation of the unexpected word itself. Additionally, the experiment explored the effect of constraint on sentence repetition effects and the downstream consequences of high constraint predictions that get violated. To investigate these issues, this pre-registered study employed a sentence repetition paradigm that manipulated the repetition of context (repetition of sentence vs repetition of word in new sentence) as well as constraint (high vs low constraint unexpected sentences).

### Sentence-level update of sematic predictions

Repeated sentences with unexpected continuations demonstrated a stronger N400 reduction from first to second presentation compared to a repetition of the previously unexpected word in a new sentence. Words that were repeated within a new sentence context could have only benefited from adaptation at the word-level. The increased repetition effect (more positive N400 upon repetition) for sentences indicates an (additional) mechanism operating at the sentence-level, updating the predictions made from the respective sentence context. While many theoretical models of predictive processing in language suggest that any unpredicted input leads to some degree of adaptation of the predictive model (Bornkessel-Schlesewsky & Schlesewsky, 2019; Fitz & Chang, 2019; Kuperberg et al., 2020; Rabovsky et al., 2018), the extend of this implicit prediction update beyond update at the word level has so far not been experimentally tested (see Besson & Kutas, 1993; Mitchell et al., 1993 for similar results from paradigms with explicit memory tasks). The current results illustrate a precise adaptation mechanism, in which unexpected semantic information influences language processing beyond the respective unexpected word and updates future predictions made from the sentence context in which the error was first experienced, possibly via adjusting connection weights between the sentence context and the previously unexpected item (Rabovsky et al., 2018). Interestingly, there was barely a repetition benefit visible for words that were repeated in a new context (the N400 seems narrower rather than reduced in amplitude). As implicit memory effects for unexpected words presented in isolation have previously been demonstrated after over 100 intervening sentences (Hodapp & Rabovsky, 2021; Rabovsky et al., 2012), the absence of a clear word repetition effect (compared to e.g., Rommers & Federmeier, 2018a, who presented only two intervening sentences) could be caused by a combination of fading implicit memory due to the amount of intervening sentences and the new context partly overwriting effects that would otherwise still be visible in a single word presentation.

In the time-frequency domain, repeated sentences were characterized by a stronger desynchronization effect in the alpha and beta band compared to the repeated presentation of the same critical words in a new sentence context. This effect has often been related to a reactivation of memory traces (Klimesch, Schack, & Sauseng, 2005). More recently, alpha/beta power decreases have been suggested to enable the processing of information (e.g. retrieval of memory traces) rather than reflecting the retrieved content itself (Griffiths et al., 2019, 2021). This interpretation of alpha/beta power changes as a more general correlate of information processing rather than a specific mental operation or content is supported by findings on the neuronal level, where suppression of alpha and beta oscillations is thought to generally increase signal-to-noise ratio by reducing the amount of synchronized spiking of neurons (Averbeck et al., 2006; Harris & Thiele, 2011). The time-frequency findings therefore provide additional evidence that the repeated sentence context can be used for easier information processing beyond the repetitions benefit of the repeated critical word itself.

### Prediction update irrespective of sentence constraint

Both, high and low constraint sentences with unexpected continuations exhibit a reduced N400 amplitude upon second encounter compared to first presentation, with no evidence for an additional benefit or disadvantage of violated high certainty predictions (only in high constraint sentences) on this reduction of ERP amplitude. In fact, Bayesian methods allowed us to find moderate evidence against such an effect (i.e., against an interaction between constraint and repetition; see Fig. 4B and 6). The overall more negative N400 for low constraint sentences at the first presentation of the sentences made the results more difficult to interpret (see Fig. 1B for a visualization of the expected pattern). In reference to the majority of the N400 literature we did not expect an effect of sentence constraint on N400 amplitudes (Federmeier et al., 2007; Kuperberg et al., 2020; Kutas & Hillyard, 1984).

However, other experiments previously did report such a difference (Federmeier & Kutas, 1999; Lai et al., 2021) and attributed it to differences in attention in strongly versus weakly constraining sentences. Another possible explanation is an overlap of components: unexpected words in strongly constraining sentences often elicit a stronger frontal late positive ERP component than in weakly constraining sentences (Brothers et al., 2020; Federmeier et al., 2007; Kuperberg et al., 2020). This frontal late positive ERP component (also sometimes referred to as post-N400-positivity: PNP) could have an earlier onset for some participants/trials and influence the overall waveform. The latter was supported by the topographical similarity of effects with both showing a left frontal maximum (Fig. 3C), which is uncommon for the N400, and an additional decomposition analysis using the RIDE toolbox (Ouyang et al., 2011). After isolating the N400 component, the effect of constraint on the first presentation was diminished (see also Fig. 5C for the corrected topographies) and statistically inconclusive, while data still showed evidence against differences in repetition effect between highly and weakly constraining sentences. Together, these findings suggest that the sentence-level prediction update when encountering unpredicted information is based on the same principles as adaptation to unexpected words themselves, as both mechanisms seem unaffected by initial sentence constraint (for word-level results see Lai et al., 2021).

The finding that unexpected input, regardless of constraint manipulations, leads to larger repetition effects (amplitude at first presentation minus second presentation) than their expected counterparts (Besson et al., 1992; Lai et al., 2021; Mitchell et al., 1993; Rommers & Federmeier, 2018b, 2018a) suggests that unexpected semantic information might be driving repetition effects. Since the focus of the current study was on the processing of unexpected information, we did not try to replicate this well-established finding and did not include an expected condition to keep the duration of the experiment manageable. However, to corroborate the general idea that a larger N400 amplitude at first encounter (presumably indicating enhanced prediction error) induces an enhanced reduction of N400 amplitude at repeated presentation (presumably reflecting enhanced adaptation), we supplement an analysis that provided strong evidence for an effect of initial N400 amplitude on the size of the N400 reduction upon repetition (repetition effect: first presentation – second presentation) (*β* = 0.94 [0.90, 0.98], BF10 = 16.09) in the sense that a larger initial N400 produced a stronger repetition induced reduction. Taken together, results from repetition paradigms could be explained by interpreting the N400 as a semantic prediction error that improves future predictions (Bornkessel-Schlesewsky & Schlesewsky, 2019; Kuperberg et al., 2020; Rabovsky et al., 2018). These theories propose that the prediction error reflected in the larger N400 amplitude for unpredicted input drives semantic adaptation, which is reflected in a larger reduction of N400 amplitude from first to second presentation. The current results add to this view by demonstrating that this reduction (i.e., adaptation) doesn’t seem to be influenced by constraint, even for sentence-level prediction adaptation. While high constraint unexpected sentences were characterized by a late frontal positive component (indexing the violation of a precise prediction), this additional cognitive process doesn’t seem to influence downstream predictive language processing at the N400 level, as amplitude reductions didn’t differ between constraint conditions. These results help establish the amount of unpredicted semantic information, rather than the precision of violated predictions, as a potential mechanism driving updated implicit predictions as reflected in reduced N400 amplitudes.

In the time-frequency domain, the onset of sentence endings showed a typical early broadband power increase and alpha and beta power decrease at their first presentation, irrespective of constraint (Rommers et al., 2017; Rommers & Federmeier, 2018b). The repetition of sentences with unexpected continuations showed a reduction in the alpha and beta band, also similar in both constraint conditions, replicating previous findings from word repetition (Burgess & Gruzelier, 2000; Klimesch et al., 1997; Rommers & Federmeier, 2018b, 2018a; Van Strien et al., 2007) in a sentence repetition paradigm. Since the alpha/beta power reduction with repetition does not seem to be influenced by sentence constraint, it can be seen as additional evidence that repeated sentences benefit from easier information processing due to previous experience irrespective of the precision of the violated predictions. We want to emphasize that the high constraint unexpected sentences still ended on a plausible continuation (but with low cloze probability), the pattern of results could thus differ for anomalous continuations (typically eliciting a semantic P600; e.g. Kuperberg et al., 2020).

### Consequences of high certainty mispredictions

While plausible high constraint prediction violations do not seem to influence the N400 amplitude at second presentation, the processing of these sentences seems to undergo additional changes, nevertheless. While the first presentation of high constraint sentences violating high certainty predictions was characterized by a late frontal positivity compared to the same unexpected item in low constraint sentences, this difference was diminished upon second presentation of the sentences. There are different interpretations of the cognitive process underlying the frontal positivity such as the update of message level representations (Brothers et al., 2023; Kuperberg et al., 2020) or the suppression of the predicted sentence continuation (Ness & Meltzer-Asscher, 2018). While future research is necessary to disentangle these accounts, the results demonstrate that learning based on just one exposure to an unexpected sentence continuation can reduce consequences of strong violated predictions.

In the time-frequency domain, our results revealed theta band effects mirroring the pattern of the frontal positivity. The analysis replicated an effect of stronger synchronization in the theta band with a fronto-central distribution for prediction violations in strongly constraining sentences compared to weakly constraining sentences that has previously been reported in the literature (Hald et al., 2006; Lewis & Bastiaansen, 2015; Pu et al., 2020; Rommers et al., 2017; Wang et al., 201). For similar paradigms, theta effects have been suggested to correlate with lexical ambiguity-resolution (Strauß et al., 2014), lexical-access (Bastiaansen et al., 2005; Hald et al., 2006) and more generally cognitive control beyond language processing (Cavanagh et al., 2010; van de Vijver et al., 2011). These explanations are not mutually exclusive, as control might be necessary to suppress the expected semantic information, activate unpredicted semantic information and/or update the current event representation (Klimesch et al., 2010; Rommers et al., 2017). The theta power band increase was only visible for highly constraining unexpected sentences and was then reduced when the sentences were repeated (i.e., leading to an interaction of constraint and repetition). Thus, the theta power effect gives additional support to the conclusions drawn from the analysis of the frontal positivity, supporting the view that the consequence of high constraint prediction violations (speculatively: additional cognitive control) is reduced after just one exposure to the unexpected sentence continuation. Subtraction of the phase-locked ERP from the signal suggested that the initial prediction violation effect on theta power seems to some extent independent of the frontal positivity ERP in a similar time window (see also Rommers et al. 2017). However, the larger reduction in theta power for prediction violations (first presentation – second presentation), as compared to unexpected words in weakly constraining contexts, was diminished after the ERP was subtracted. It is therefore likely that the underlying processes are at least partly shared/overlapping, suggesting a connection worth investigating in future work.

In general, it seems not entirely clear whether additional adaptation has taken place for high constraint unexpected sentences, or whether the same adaptation happened in the high and low constraint sentences but with partly different consequences. Additional adaptation could mean that in high constraint sentences, a strong violation leads to an additional update that is reflected not in N400 amplitudes but in the other measures such as the frontal positivity and theta power. The same adaptation but with different consequences could mean that in the high constraint sentences, adaptation might have reduced the need for suppression of the predicted sentence continuation, update of higher-level representations and/or cognitive control (processes that seem highly compatible), possibly because after the adaptation in response to unexpected input, there was less of a strong pre-activation of a specific continuation. The consequence of the same adaptation for low constraint sentences might have been more gradual because there was no strong specific pre-activation to begin with, even before the adaptation. We prefer the latter interpretation (same adaptation, different consequences), because it seems more in line with the common interpretations of the involved measures (the N400, the late frontal positivity, and theta power), but future work would be desirable to further disentangle these possibilities.

### Explicit memory effects

There were no LPC effects neither for the context nor constraint manipulation analysis. Even though LPC effects are thought to persist longer than N400 effects of repetitions, the higher number of intervening sentences could still explain the absence of an LPC effect for words repeated in new sentences. However, for repeated sentences LPC effects have been demonstrated after a lag of up to 45 min (Besson et al., 1992). One possible explanation for the current result is that participants didn’t engage in deep processing when reading the sentences. It has been repeatedly shown that while the N400 seems to reflect an implicit process that doesn’t require such engagement and newer results suggest the same for the frontal positivity (Lai et al., 2023), LPC effects are strongly influenced by the depth of processing at encoding (Paller et al., 1995; Paller & Kutas, 1992; Rugg et al., 1998). Since LPC effects are usually seen as an index for explicit memory and conscious recollection (Smith, 1993; Van Petten & Senkfor, 1996; Wilding et al., 1995; Wilding & Rugg, 1996), the absence of these effects additionally suggests that participants didn’t consciously remember or recognize the repeated material. This could be seen as an indicator that explicit memory played little to no role while processing the sentences. The declarative memory literature suggests that results might differ if sentences were processed more deeply. In the memory and schema literature there is evidence for an explicit memory benefit for schema congruent information (potentially via the ventromedial prefrontal cortex, inhibition of medial temporal lobe (MTL) and the strengthening of neocortical connections) but also incongruent or novel experiences (potentially via MTL and the encoding of a specific instance; Gilboa & Marlatte, 2017; Greve et al., 2019; Van Kesteren et al., 2012). A similar sentence paradigm as the one employed here, but with an explicit learning task, revealed better memory for expected sentence endings compared to unexpected sentence endings and a general explicit memory benefit for information presented in strongly constraining sentences (Höltje & Mecklinger, 2022). Based on the explicit memory literature focusing specifically on the processing of unexpected information, a benefit in declarative learning for unexpected information violating a high certainty prediction would be expected (Gambi et al., 2021; Greve et al., 2017), potentially by attracting more attention (Butterfield & Metcalfe, 2006). While we were interested in implicit language adaptation and LPC results support a reduced role of explicit memory effects, subject or sentence-level differences in the contribution of the declarative memory system could potentially explain the only moderate evidence against an effect of the violated prediction’s precision on adaptation. Disentangling the contributions of different memory systems to predictive language processing and model update is therefore an exciting future perspective.

### No evidence for pre-stimulus differences depending on constraint

We found no evidence for a pre-stimulus difference between high constraint and low constraint sentences, even though this comparison has previously been characterized by differences in the alpha or beta band (Gastaldon et al., 2020; León-Cabrera et al., 2022; Li et al., 2017; Piai et al., 2014; Rommers et al., 2017; Wang et al., 2018). Terporten et al. (2019) suggested that pre-stimulus alpha and beta desynchronization was not monotonically related to constraint and therefore does not seem to be directly reflective of predictive processes.

Instead, the authors found the strongest decrease for intermediately constraining sentence contexts (as compared to strongly constraining sentences as used here), which might explain the absence of effects in the current study (see also e.g., Rommers & Federmeier, 2018a).

## Conclusions

Together, the results presented here suggest a predictive language system that is constantly and quickly adapting to the statistics of the environment. One characteristic that makes this predictive mechanism so effective is that a single encounter with unpredicted semantic information can already update predictions made from the same sentence context and facilitate information processing, beyond just changing the representations of the respective unexpected word itself. On the level of the N400 ERP component and the alpha/beta frequency bands, this one-shot adaptation of sentence-based predictions lead to the same degree (neither more nor less) of facilitated processing in high and low constraint sentences with unexpected endings, supporting previous findings that the precision of violated predictions does not seem to be critical to the implicit update of semantic predictions.

Combining the current results on constraint in unexpected sentence repetition with previous findings in literature that have repeatedly shown a larger reduction of N400 amplitudes upon repetition for unexpected sentence continuation compared to predictable continuations, this pattern would be in line with theories proposing that unpredicted semantic information (as reflected in the N400) is driving adaptation effects. Additionally, adaptation in predictive sentence processing has consequences beyond semantic prediction update. While high constraint unexpected sentence continuations seem to require some form of cognitive control to deal with strong mispredictions, the flexible adaptation of the language model eliminates the necessity of these additional resources, if the same context is processed again.

## Data availability statement

The data used for generating the figures and analysis scripts are openly accessible at: https://osf.io/rk2du/. The raw EEG data cannot be made available in a public repository due to the privacy policies for human biometric data according to the European General Data Protection Regulation (GDPR). Further preprocessed data can be obtained from the corresponding author upon reasonable request and as far as the applicable data privacy regulations allow it.

## Author contribution statement

Conceptualization: AH, MR; Methodology, Visualization, formal Analysis: AH supervised by MR; Writing - Original Draft Preparation: AH; Writing - Review & Editing: MR; Supervision: MR; Funding Acquisition: MR.

## Acknowledgements

We thank Jessica Hochwald, Nina Koch, and Lioba Berndt for their help with data collection and/or stimulus preparation. Funding was provided by an Emmy Noether grant from the German Research Foundation (grant RA 2715/2-1) to Milena Rabovsky.

## Footnotes

1 This means that we did not assess the size of repetition effects in our manipulations in comparison to expected sentence continuations. However, the effect of larger repetition effects for unexpected compared to expected sentences is well-established in the previous literature (e.g., Besson et al., 1992; Lai et al., 2021; Mitchell et al., 1993; Rommers & Federmeier, 2018b, 2018a). The current study therefore focuses only on understanding the repetition effect for unexpected continuations.

2 Approximate dimensions of the clusters are provided as informative descriptions and not inferential claims about onset and offset of time window, spatial distribution, or frequencies. Cluster based permutation tests only control the false alarm rate under the null hypothesis that there is no effect between the conditions (Maris & Oostenveld, 2007; Piai et al., 2015; Sassenhagen & Draschkow, 2019).

